# DeepMAPS: Single-cell biological network inference using heterogeneous graph transformer

**DOI:** 10.1101/2021.10.31.466658

**Authors:** Anjun Ma, Xiaoying Wang, Cankun Wang, Jingxian Li, Tong Xiao, Juexing Wang, Yang Li, Yuntao Liu, Yuzhou Chang, Duolin Wang, Yuexu Jiang, Jinpu Li, Li Su, Shaopeng Gu, Gang Xin, Zihai Li, Bingqiang Liu, Dong Xu, Qin Ma

**Author notes:** These authors contributed equally to the paper as first authors (Anjun Ma, Xiaoying Wang). To whom correspondence should be addressed, **Corresponding author**, Correspondence to Dr. Qin Ma:; Dr. Dong Xu; Dr. Bingqiang Liu.

## Abstract

We present DeepMAPS (Deep learning-based Multi-omics Analysis Platform for Single-cell data) for biological network inference from single-cell multi-omics (scMulti-omics). DeepMAPS includes both cells and genes in a heterogeneous graph to simultaneously infer cell-cell, cell-gene, and gene-gene relations. The multi-head attention mechanism in a graph transformer considers the heterogeneous relation among cells and genes within both local and global context, making DeepMAPS robust to data noise and scale. We benchmarked DeepMAPS on 18 scMulti-omics datasets for cell clustering and biological network inference, and the results showed that our method outperformed various existing tools. We further applied DeepMAPS on lung tumor leukocyte CITE-seq data and matched diffuse small lymphocytic lymphoma scRNA-seq and scATAC-seq data. In both cases, DeepMAPS showed competitive performance in cell clustering and predicted biologically meaningful cell-cell communication pathways based on the inferred gene networks. Note that we deployed a webserver using DeepMAPS implementation equipped with multiple functions and visualizations to improve the feasibility and reproducibility of scMulti-omics data analysis. Overall, DeepMAPS represents a heterogeneous graph transformer for single-cell study and may benefit the use of scMulti-omics data in various biological systems.

## Main

Single-cell sequencing, such as single-cell RNA sequencing (scRNA-seq) and single-cell ATAC sequencing (scATAC-seq), reshapes the investigation of cellular heterogeneity and yields novel insights in neuroscience, cancer biology, immuno-oncology, and therapeutic responsiveness^1, 2^. However, an individual single-cell modality only reflects a snapshot of genetic features and partially depicts the peculiarity of cells, leading to characterization biases in complex biological systems^2, 3^. Single-cell multi-omics (scMulti-omics) allows for the generation and quantification of multiple modalities simultaneously to fully capture the intricacy of complex molecular mechanisms and cellular heterogeneity. Such analyses advance various biological studies when paired with robust computational analysis methods^4^.

The existing tools for integrative analyses of scMulti-omics data, e.g., Seurat^5^, MOFA+^6^, Harmony^7^, and totalVI^8^, reliably predict cell types and states, remove batch effects, and reveal relationships or alignments among various modalities. However, most existing methods do not explicitly consider in their models the crosstalk between cells or different molecular modalities. Hence, these models cannot infer the active biological networks of diverse cell types and have limited power to elucidate the response of these complex networks to external stimuli in specific cell types.

Recently, graph neural networks (GNN) have shown a unique strength in learning low-dimensional representations of individual cells by propagating neighbor cell features and constructing cell-cell relations in a global cell graph^9–12^. For example, our in-house tool scGNN, a novel GNN model, has demonstrated superior cell clustering and gene imputation performance based on large-scale scRNA-seq data^13^. Furthermore, a heterogeneous graph with different types of nodes and edges has been widely used to model a multi-relational knowledge graph^14^. It provides a natural representation framework for integrating scMulti-omics data and learning the underlying cell-type-specific biological networks. Moreover, a recent development in the attention mechanism for modeling and integrating heterogeneous relationships has made deep learning models explainable and enabled the inference of cell-type-specific biological networks^14, 15^.

To this end, we developed **DeepMAPS** (**Deep** learning-based **M**ulti-omics **A**nalysis **P**latform for **S**ingle-cell data), a heterogeneous graph transformer framework for cell-type-specific biological network inference from scMulti-omics data. DeepMAPS formulates high-level representations of relations between cells and genes in a heterogeneous graph, with cells and genes as the two disjoint node sets in this graph. Projecting the features of genes and cells into the same latent space is an effective way to harmonize the imbalance of different batches and create a solid foundation for cell clustering (i.e., node clustering) as well as the prediction of cell-gene and gene-gene relations in a specific cell cluster (i.e., link prediction). Most importantly, the attention mechanism in this transformer model enhances biological interpretability and enables the identification of active gene modules in each cell cluster. Overall, DeepMAPS is an end-to-end and hypotheses-free framework that provides the first deep learning tool to infer cell-type-specific biological networks from scMulti-omics data. We deployed DeepMAPS as a code-free web portal along with Docker to ensure the reproducibility of scMulti-omics data analysis and lessen the programming burden for biologists who lack sufficient computational skills or resources (**Fig. 1**).

**Fig. 1.**
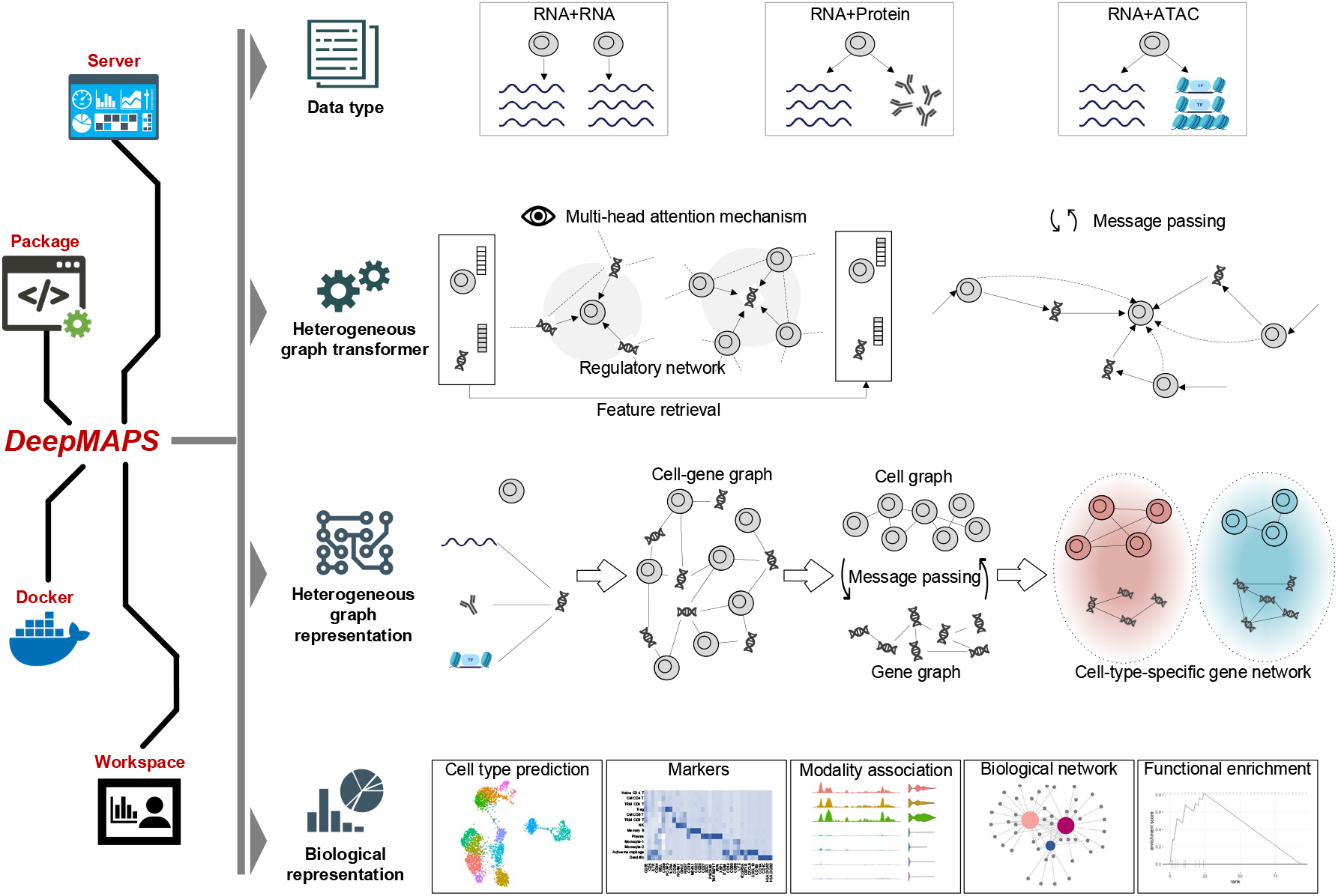
DeepMAPS is a Deep learning-based Multi-omics Analysis Platform for Single-cell data. It allows for the joint analysis of multiple scRNA-seq, CITE-seq (matched RNA and protein profilings), and matched single-cell RNA and ATAC-seq data sets. The core method includes the representation of cell-gene relations via a heterogeneous graph and a transformer with a graph attention mechanism. DeepMAPS provides interactive and interpretable graphical representations to deliver cell clusters and various cell-type-specific biological networks based on modality types. DeepMAPS is provided as a web portal to ensure robustness and reproducibility, along with a docker container. Workspace is provided for job saving and retrieval. DeepMAPS also supports diverse interpretations, including but not limited to joint cell clustering, marker identification, modality associations, cell-type-specific biological network inference, and functional enrichment.

## Results

### Overview of DeepMAPS

There are five steps in the DeepMAPS pipeline to accomplish a joint analysis of scMulti-omics data (**Fig. 2** and **Methods**). (*i*) Data are preprocessed by removing low-quality cells and modalities,= and then different normalization methods are applied according to the specific data types. (*ii*) An integrated cell-gene matrix is generated to represent the combined activity of each gene in each cell. Different data integration methods are applied for different scMulti-omics data types. (*iii*) A heterogeneous graph transformer (**HGT**) model is built to jointly learn the low-dimensional embedding for cells and genes and generate an attention score to indicate the importance of a gene to a cell. (*iv*) Cell clustering and functional gene modules are predicted based on HGT-learned embeddings and attention scores. (*v*) Diverse biological networks, e.g., gene regulatory networks and gene association networks, are inferred for each cell type.

**Fig. 2.**
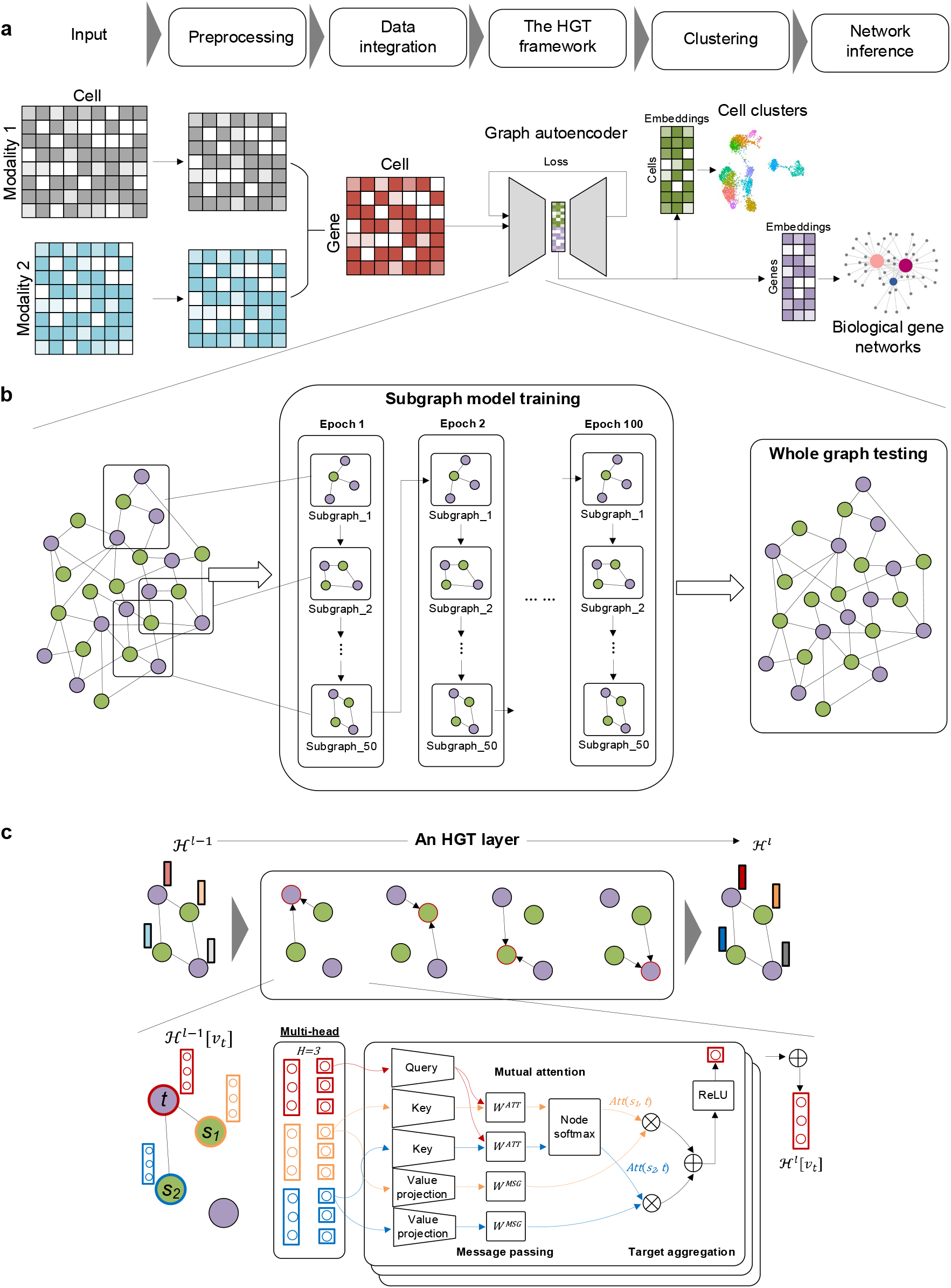
DeepMAPS framework. (**a**) The five major steps and graphical illustration of DeepMAPS. (**b**) Relations in the heterogeneous graph of cells and genes are learned in an HGT-based graph autoencoder. The hyperparameters are first trained in 50 subgraphs and 100 epochs and then applied to test the whole graph. (**c**) For each subgraph training and whole graph testing, multiple HGT layers are applied. An example with two cells (purple) and two genes (green) is shown for one HGT layer. The embeddings of the target cell (red) and two neighbor genes (orange and blue) are separated evenly into three heads. For each head, the HGT layer calculates the attention score of neighbor genes to the target cell and updates the target’s embedding of this head. The entire target cell embedding is updated by concatenating all three heads to complete one HGT layer (Method).

To learn joint representations of cells and genes, we first generate a cell-gene matrix integrating all heterogeneous information of the input scMulti-omics data. A heterogeneous graph with cell nodes and gene nodes is then constructed, wherein a cell-gene edge represents the integrated gene activity score in the matrix, and the initial embedding of each node is learned from the gene-cell integrated matrix via a two-layer GNN graph autoencoder. The entire heterogeneous graph is then sent to a graph autoencoder to learn relations between the cells and genes and update the embedding of each node. Here, DeepMAPS adopts a heterogeneous multi-head attention mechanism to model the overall topological information (global relationships) and neighbor message passing (local relationships) on the heterogeneous graph.

In each layer, each node (either a cell or a gene) is considered a target and its 1-hop neighbors as sources. DeepMAPS evaluates the importance of its neighbor nodes and the amount of information that can be passed to the target based on the synergy of node embedding. As a result, cells and genes with high positively correlated embeddings are more likely to pass messages within each other, thus maximizing the similarity and disagreement of the embeddings. To make the unsupervised training process feasible on a large heterogeneous graph, DeepMAPS is first performed on 50 subgraphs sampled from the heterogeneous graph, covering a minimum of 50% of all nodes to train for the shared parameters between different nodes, information which is later used for testing of the whole graph. As an important training outcome, an attention score is given to represent the importance of a gene to a cell. A high attention score for a gene to a cell implies that the gene is relatively important for defining cell identity and characterizing cell heterogeneity. This discrimination allows for the construction of reliable gene association networks in each cell cluster as the final output of DeepMAPS. We then build a Steiner Forest Problem (SFP) model^16^ using to identify genes with higher attention scores and similar embedding features unique to a cell cluster. The gene-gene and gene-cell relations in the optimized solution of the SFP model mirror the co-expression relations between genes and the specificity of gene attention to a cell type. A gene network established from SFP contains genes that are of the most important in characterizing the identity of that cell cluster based on their attention scores and embedding similarities, and these genes are considered to be cell-type-active.

### DeepMAPS achieves superior performances in joint cell clustering and biological network inference from scMulti-omics data

We evaluated DeepMAPS on 18 scMulti-omics datasets, including two multiple scRNA-seq data sets (Data 1-2), eight CITE-seq datasets (Data 3-10), and eight matched scRNA-seq and scATAC-seq (scRNA-ATAC-seq) datasets measured from the same cell (Data 11-18) (**Supplementary Data 1**). Specifically, Data 1–2 and 17–18 have benchmark annotations provided in the original manuscripts. These data cover a number of cells ranging from 549 to 30,672; an average read depth (considering scRNA-seq data only) ranging from 2,933 to 645,526; and a zero-expression rate (considering scRNA-seq data only) from 71% to 97% (**Fig. 3a**).

**Fig. 3.**
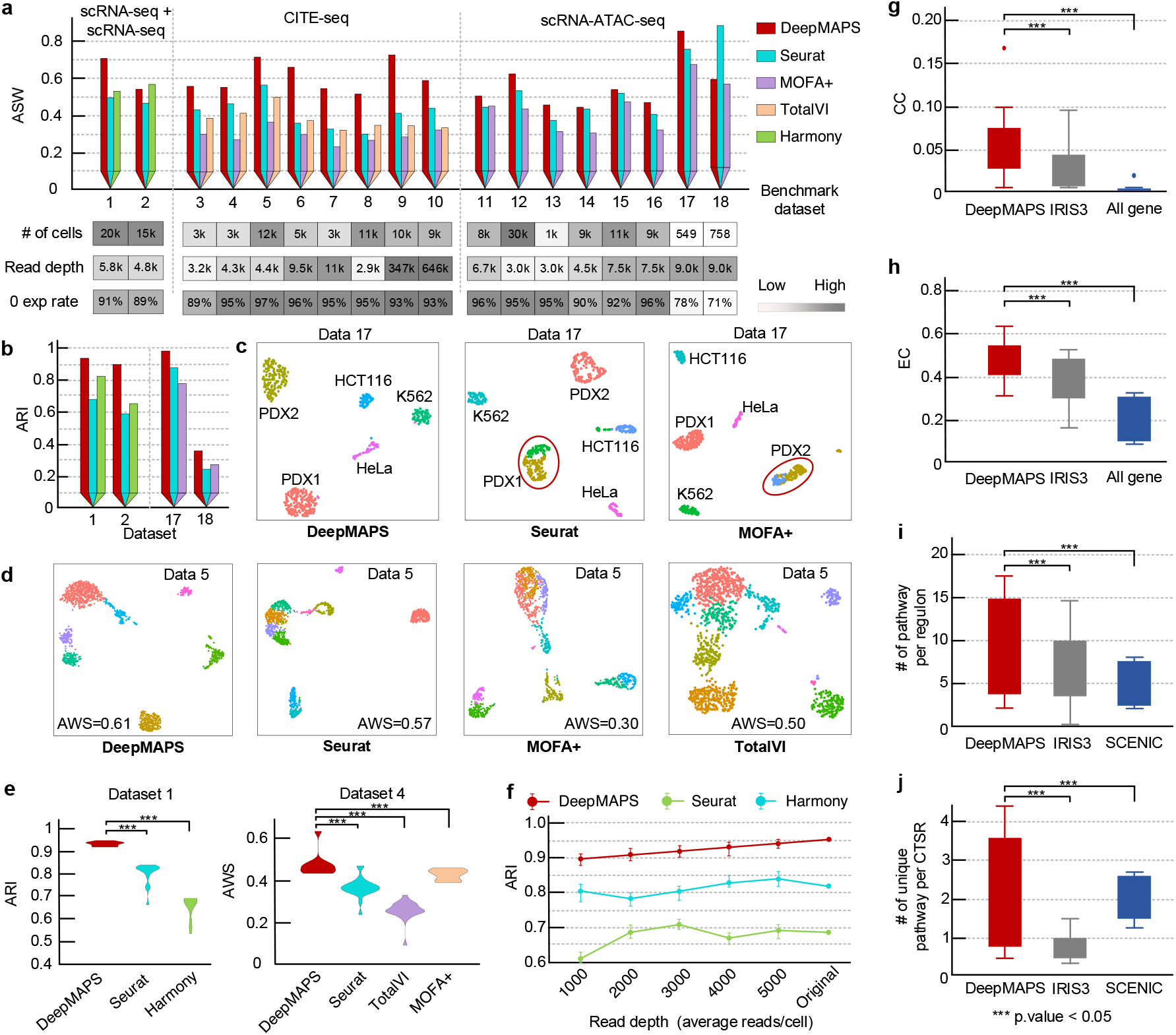
Benchmarking of DeepMAPS in cell clustering and biological network inference. (**a**) Benchmark cell clustering results of all 18 datasets in terms of ASW without using benchmark cell labels. Different benchmarking tools were selected for comparison based on the capability of the tool. The heatmaps indicate the number of total cells, average gene expression read depth per cell, and average RNA zero-expression rate in each data. (**b**) Results comparison on four datasets with benchmarking cell labels in terms of adjusted rand index. (**c**) UMAP comparison of Data 17 (with benchmark labels) between DeepMAPS and other tools. Cluster labels were annotated based on cell correspondence to the original cell label. (**d**) UMAP comparison of Data 5 (without benchmark labels) between DeepMAPS and other tools. (**e**) Robustness test of DeepMAPS using the cell cluster leave-out method for Data 1 (with benchmark labels) and Data 4 (without benchmark labels). Details can be found in the Method section. (**f**) Robustness test of DeepMAPS to different read depths with Data 1. (**g** and **h**) Evaluation and comparison of gene association network inference of DeepMAPS and other methods. Closeness centrality and betweenness centrality were used to indicate the compactness and connectivity of networks inferred from different methods. (**i** and **j**) Evaluation and comparison of gene regulatory networks (regulons) identified by DeepMAPS, IRIS3, and SCENIC, based on the number of functional pathways enriched in a regulon or cell-type-specific regulon.

We compared DeepMAPS with four benchmarking tools (Seurat, MOFA+, TotalVI, and Harmony (**Methods**)) using the default settings in terms of the Average Silhouette Width (ASW) (**Fig. 3a**) and Adjusted Rand Index (ARI) (**Fig. 3b**) to evaluate the performance of joint cell clustering for all three scMulti-omics data types. DeepMAPS was trained by each scMulti-omics data type and each dataset in an unsupervised way. One set of parameters was chosen as the default for all datasets within the same data type based on the grid optimization of hyper-parameter combinations (**Supplementary Data 2-4**). The results showed that, in all three scMulti-omics data scenarios, DeepMAPS distinctly outperformed the other tools in most cases. Note that for Data 2 and 18, though DeepMAPS did not achieve the best ASW compared to the other tools, its performance regarding ARI comparison to the benchmark label was the highest.

We show the UMAP results for the cell clustering of Data 17, a cancer cell line (n=549) with benchmarked cell labels for scRNA-ATAC-seq data (**Fig. 3c**). Compared with the original cell line labels, DeepMAPS was the only tool that accurately separated each cell type with minimum mismatches (ARI=0.97). In contrast, Seurat (ARI=0.88) and MOFA+ (ARI=0.79) mistakenly divided the PDX1 or PDX2 population into two clusters and included more mismatches. DeepMAPS also produced better UMAP visualization than other tools for datasets without benchmark labels, without mixing clusters or separating random cells (Fig. 3d).

In order to evaluate the robustness of DeepMAPS, we performed a leave-out test on benchmark datasets (**Supplementary Data 5**). For data with benchmark labels, we first filtered cells by removing a cluster of cells based on benchmark labels and then performed analysis using DeepMAPS and other tools; for other datasets, we removed cells based on clusters identified by each tool. The results showed that DeepMAPS achieved better performance compared to all other benchmarking methods (**Fig. 3e)**. Another test was performed on a series of data simulated for different read depth rates (**Fig. 3f**, **Supplementary Data 6**). For each testing dataset, the clustering results of DeepMAPS were consistent with high AWS or ARI scores, indicating that the message passing and attention mechanism used by DeepMAPS helped maintain cell-cell relations and tolerance to varying read depths.

We further evaluated the two kinds of biological networks that DeepMAPS can produce (**Supplementary Data 7**). For the gene association network for all scMulti-omics data types, we used the centrality scores and enriched pathways to compare DeepMAPS with IRIS3^11^. IRIS3 is an in-house tool for identifying cell-type-specific regulons and constructing gene regulatory networks from scRNA-seq data. It has superior performance over other public tools, such as SCENIC^10^. We also compared our results to the co-expression network constructed from the whole gene list with significance cutoffs, which is widely used in single-cell studies. Both the average closeness centrality and eigenvector centrality scores of networks constructed by DeepMAPS showed significantly higher scores than the other two methods when using all 18 benchmark datasets (**Fig. 3g-h**). Moreover, for the gene regulatory network constructed from scRNA-ATAC-seq data, we evaluated the number of significantly enriched pathways in a TF-regulon and a cell-type-specific regulon (**Fig. 3i-j**). The results indicated that DeepMAPS could construct more compatible and biologically reasonable gene networks for each cell type and outperform the other methods.

### DeepMAPS accurately identify cell types and infer cell-cell communication in PBMC and lung tumor immune CITE-seq data

We present a case study that applies DeepMAPS to a published mixed peripheral blood mononuclear cells (PBMC) and lung tumor leukocytes CITE-seq dataset to demonstrate capacity in modeling scMulti-omics in characterizing cell identities. The dataset includes RNAs and proteins measured on 3,485 cells. DeepMAPS identified 13 cell clusters, including four CD4^+^ T cell groups (naïve, central memory (CM), tissue-resident memory (TRM), and regulatory (Treg)), two CD8^+^ T cell groups (CM and TRM), a natural killer cell group, a memory B cell group, a plasma cell group, two monocyte groups, one tumor-associated macrophage (TAM) group, and a dendritic cell (DC) group. We annotated each cluster by visualizing the expression levels of curated maker genes and proteins (**Fig. 4a-b**, **Supplementary Data 8**). Compared to cell types identified using only proteins or RNA, we isolated or accurately annotated cell populations that could not be characterized using the individual modality analysis. In summary, by combining signals captured from both RNA and proteins, DeepMAPS successfully identified biologically reasonable and meaningful cell types in the CITE-seq data.

**Fig. 4.**
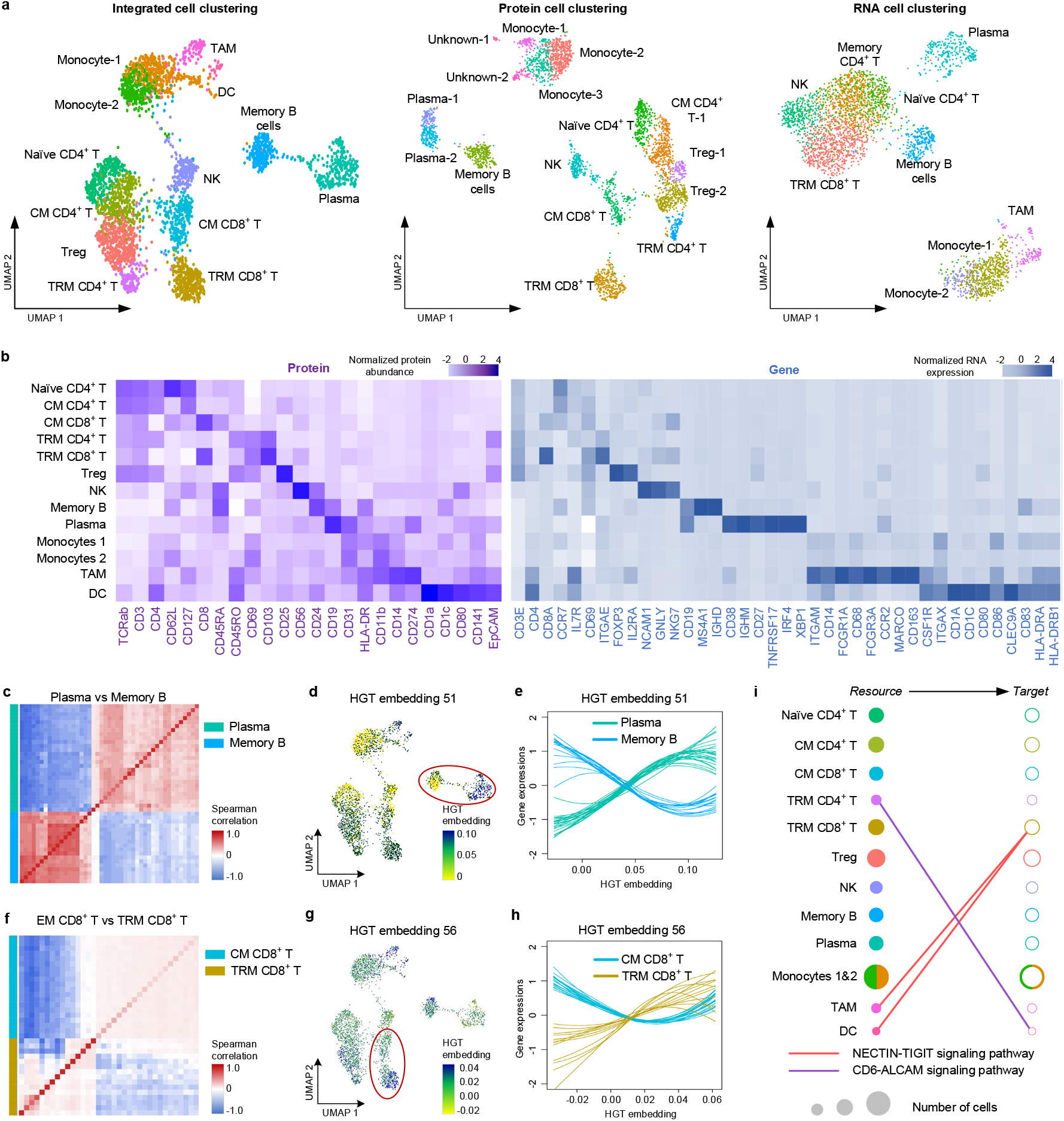
DeepMAPS identification of heterogeneity in CITE-seq data of PBMC and lung tumor leukocytes. (**a**) UMAPs for DeepMAPS cell clustering results from integrated RNA and protein data, protein data only, and RNA data only. Cell clusters were annotated based on curated marker proteins and genes. (**b**) Heatmap of curated marker proteins and genes that determine the cell clustering and annotation. (**c**) Heatmap of the correlation comparison of top differentially expressed genes and proteins in plasma cells and memory B cells. (**d**) UMAP is colored by the 51^st^ embedding, indicating distinct embedding representations in plasma cells and memory B cells. (**e**) Expression of top differentially expressed genes and proteins in ***c*** as a function of the 51^st^ embedding to observe the pattern relations between plasma cells and memory B cells. Each line represents a gene/protein, colored by cell types. For each gene, a line was drawn using a loess smoothing function based on the corresponding embedding and scaled gene expression in a cell. (**f-h**) Similar visualization was conducted to compare EM CD8^+^ T cells and TRM CD8^+^ T cells (***c-e)***. (**i**) Two signaling pathways, NECTIN and ALCAM, are shown to indicate the predicted cell-cell communications between two cell clusters. A link between a filled circle (resource cluster with highly expressed ligand coding genes) and an unfilled circle (target cluster with highly expressed receptor coding genes) indicates the potential cell-cell communication of a signaling pathway. Circle colors represent different cell clusters, and the size represents the number of cells. The two monocyte groups were merged.

We then compared the modality correlation between two cell types. We used the top differentially expressed genes and proteins between memory B cells and plasma cells and performed hierarchical clustering of the correlation matrix. The result clearly stratified these features into two anticorrelated modules: one associated with memory B cells and the other with plasma cells (**Fig. 4c**). Furthermore, we found that the features in the two modules significantly correlated with the axis of maturation captured by our HGT embeddings (**Supplementary Fig. 1**). For example, we observed that one HGT embedding (the 51^st^) showed distinctive differences between plasma cells and memory B cells (**Fig. 4d-e**). Similar findings were also observed when comparing EM CD8^+^ T cells and TRM CD8^+^ T cells and showed a much closer relationship for expression correlations (**Fig. 4f**). Nevertheless, it is possible to identify a representative HGT embedding (56^th^) that maintains embedding signals for a defined separation of the two groups (**Fig. 4g-h**). These results point to any two cell clusters consisting of coordinated activation and repression of multiple genes and proteins, leading to a gradual transition in cell state that can be captured by a specific dimension of the DeepMAPS latent HGT space. On the other hand, we generated the gene-associated networks with genes showing high attention scores for EM CD8^+^ T cells, TRM CD8^+^ T cells, memory B cells, and plasma cells and observed diverse patterns (**Supplementary Fig. 2**).

Based on the cell types and raw data of gene and protein expressions, we inferred cell-cell communication using CellChat^17^. We constructed communication networks among different cell types within multiple signaling pathways (**Fig. 4i)** and further applied CellChat^17^ to detect any ligand-receptor interactions that have been validated between any cell types we identified. For example, we observed a CD6-ALCAM signaling pathway existing between DC (source) and TRM CD4^+^ T cells (target) in the lung cancer tumor microenvironment (TME). Previous studies have shown that ALCAM on antigen-presenting DCs interact with CD6 on the T cell surface and contribute to T cell activation and proliferation^18–20^. As another example, we identified the involvement of the NECTIN-TIGIT signaling pathway during the interaction between the TAM (source) and TRM CD8^+^ T cells (target), which is supported with a previous report that NECTIN (CD155) expressed on TAM could be immunosuppressive when interacting with surface receptors, TIGIT, on CD8^+^ T cells in the lung cancer TME^21, 22^.

### DeepMAPS identifies specific gene regulatory networks in diffuse small lymphocytic lymphoma scRNA-seq and scATAC-seq data

To further extend the power of DeepMAPS to gene regulatory network inference, we used a single-cell multiome ATAC+Gene expression dataset available on the 10X Genomics website (10X Genomics online resource, **Supplementary Data 1**). The raw data is derived from 14,566 cells of flash-frozen intra-abdominal lymph node tumor from a patient diagnosed with diffuse small lymphocytic lymphoma (DSLL) of the lymph node lymph. We developed a new method to achieve the integration of gene expression and chromatin accessibility by balancing the weight of each modality of a gene in a cell, based on RNA velocity (**Fig. 5a** and **Method**). We demonstrate that using velocity-weighted integration methods can achieve better and rigorous cell clustering results than the Seurat weighted nearest neighbor method (**Supplementary Fig. 3**). To build TF-gene linkages, we considered the gene expression, gene accessibility, TF-motif binding affinity, peak to gene distance, and TF-coding gene expression. Genes found to be regulated by the same TF in a cell cluster are grouped as a regulon. We considered regulons with higher centrality scores to have greater influences on the characterization of the cell cluster. Regulons regulated by the same TF across different cell clusters are compared for differential regulon activities. Those with significantly higher regulon activity scores (RAS) are considered to be the cell-type-specific regulons in the cell cluster.

**Fig. 5.**
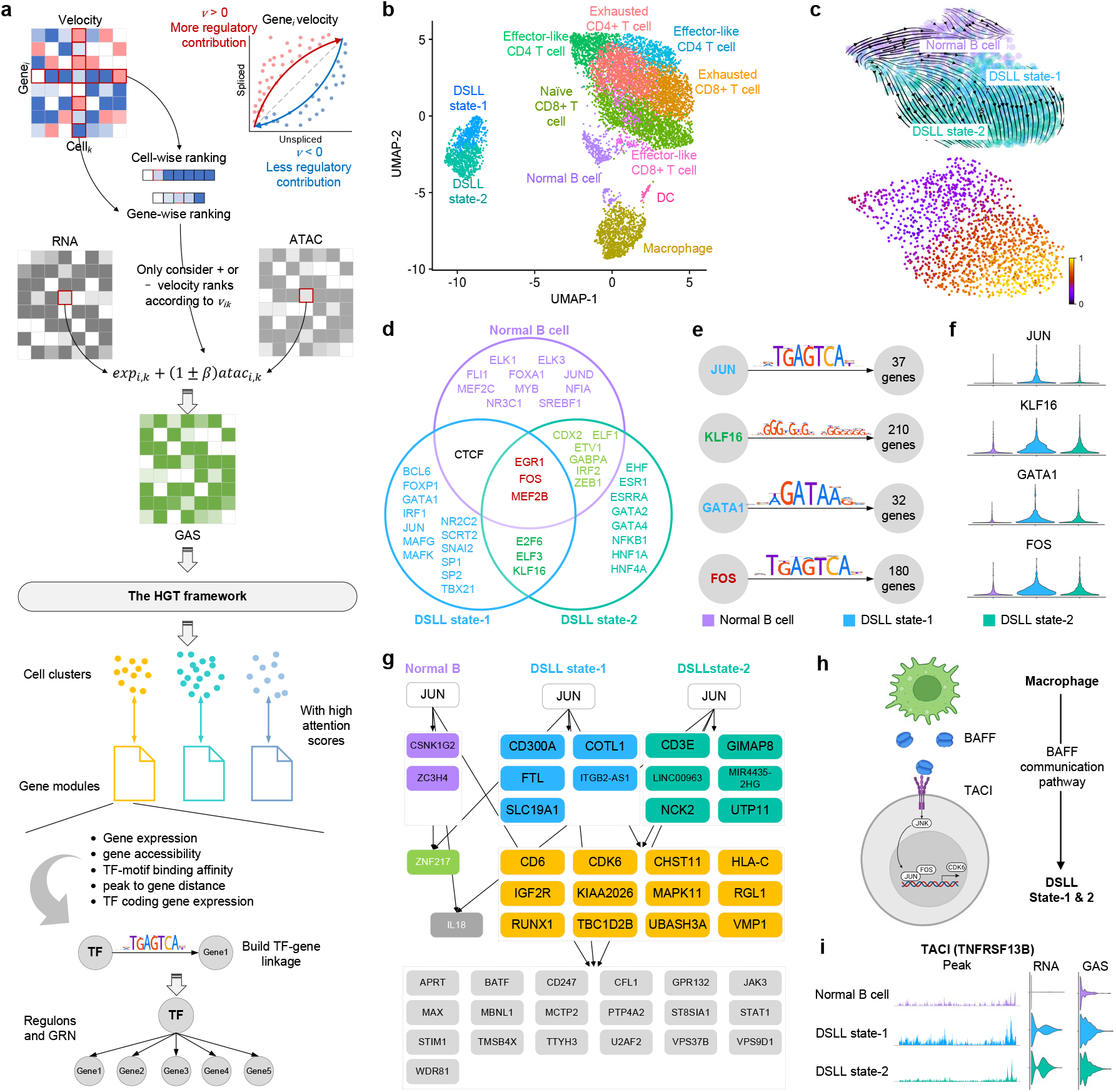
DeepMAPS identifies specific gene regulatory networks in DSLL subnetworks. (a) Conceptual illustration of DeepMAPS analysis of scRNA-ATAC-seq data. Modalities are first integrated based on a velocity-weighted balance. The integrated gene activity matrix (GAS) was then used to build a heterogeneous graph as input into the HGT framework. The cell cluster and gene modules with high attention scores were then used for building TF-gene linkages and determining regulons in each cell cluster. (b) The UMAP shows the clustering results of DeepMAPS. Cell clusters were manually annotated based on curated marker genes. (c) The observed and the extrapolated future states (arrows) based on the RNA velocity of the normal B cell and the two DSLL states are shown (top panel). Velocity-based trajectory analysis shows the pseudotime from the top to the bottom right (bottom panel). (d) Selected 20 TF in each of the three clusters, representing the top 20 regulons with the highest centrality scores. Colors represent regulons uniquely identified in each cluster or shared between different clusters. (e) Regulons in DSLL state-1 showed a significant difference in regulon activity compared to the other clusters. Motif shape and number of regulated genes are also shown. (f) Violin plots of regulon activities of the four regulons compared between the three clusters. (g) The downstream-regulated genes of JUN (the most differentially active regulon in DSLL state-1) in the three clusters. (h) An illustration of the BAFF signaling pathway identified from GAS-based cell-cell communication prediction using CellChat. The BAFF signaling pathway was found to exist between macrophage and both DSLL states. It further activates the JUN regulon and enables the transcription of genes like CDK6. (i) The ATAC peak, RNA expression, and GAS level of TNFRSF13B (the coding gene of TACI, the receptor in the BAFF signaling pathway).

DeepMAPS identified 11 cell clusters in the DSLL data. All clusters were manually annotated based on curated gene markers (**Fig. 5b**, **Supplementary Data 9**). Two DSLL-like cell clusters (DSLL state-1 and state-2) were observed. The RNA velocity-based pseudotime analysis performed on the three B cell clusters (normal B cell and two DSLL states) assumed that the two DSLL states were derived from normal B cells, and state-1 is derived earlier than state-2, although the two states seemed to be partially mixed (**Fig. 5c**). We further selected the top 20 TFs with the highest regulon centrality scores in each of the three cell clusters (**Fig. 5d**, **Supplementary Data 10**). Interestingly, these TFs showed distinctions between the normal and the two DSLL states and inferred variant regulatory patterns within the two DSLL states. For regulons shared by all three B cell clusters, EGR1, MEF2B, and FOS were transcriptionally active in both normal B and DSLL cells and responsible for regulating B cell development, proliferation, and germinal center formation^23–26^. E2F6, ELF3, and KLF16 were identified as shared only in the two DSLL states, with reported roles in tumorigenesis^27–32^. Further, JUN, MAFK, and MAFG, which encode the compartments of the activating protein-1 (AP-1),^26, 33, 34^ were found to be active in DSLL state-1 while NFKB1, coding for a subunit of the NF-κB protein complex^35, 36^, was found to be active in DSLL state-2.

We constructed a gene regulatory network consisting of the four cell-type-specific regulons (JUN, KLF16, GATA1, and FOS) (**Fig. 5e**, **Supplementary Fig. 4**) in DSLL state-1 with RAS that are significantly higher than normal B cells and DSLL state-2 (**Fig. 5f**). KLF16 reportedly promotes the proliferation of both prostate^31^ and gastric cancer cells^32^. FOS and JUN are transcription factors in the AP-1 family, regulating the oncogenesis of multiple types of lymphomas^26, 33, 34, 37^, and GATA1 is essential for hematopoiesis, the dysregulation of which is implicated in multiple hematologic disorders and malignancies^38, 39^. Distinct regulatory patterns were also observed when we zoomed in to a single regulon (**Fig. 5g**, **Supplementary Fig. 5**). As the most active regulon in DSLL state-1, JUN was found to regulate five unique downstream genes and 12 genes shared with DSLL state-2. Downstream genes, including CDK6^25, 26^, IGF2R^40^, and RUNX1^41^, are critical for cell proliferation, survival, and development functions in DSLL.

Moreover, we further built connections between upstream cell-cell communication signaling pathways and downstream regulatory mechanisms in DSLL cells. We identified a cell-cell communication between macrophage and the two DSLL states via the B cell activation factor (BAFF) signaling pathway, based on the integrated GAS matrix using CellChat^17^, which includes BAFF as the ligand on macrophage cells and TACI (transmembrane activator and calcium-modulator and cyclophilin ligand interactor) as the receptor on DSLL cells (**Fig. 5h**). BAFF signaling is critical to the survival and maturation of normal B cells^42, 43^, while aberrations contribute to the resistance of malignant B cells to apoptosis^44^, ^45^. We observed that the expression of the TACI coding gene, TNFRSF13B, was explicitly higher in the two DSLL states, while the corresponding chromatin accessibility maintained high peaks in state-1 (**Fig. 5i**). Upon engagement with its ligand, TACI has been reported to transduce the signal and eventually activate the AP-1^46, 47^ and NF-κB^48, 49^ transcriptional complexes for downstream signaling in B cells. JUN (a subunit of AP-1) was identified as the most specific and key regulator in state-1 responsible for cell proliferation and regulating downstream oncogenes, such as CDK6 that has been reported to promote the proliferation of cancer cells in multiple types of DSLLs as well as other hematological malignancies^50–52^. It is now clear that BAFF signaling first appears in DSLL state-1 and triggers the activation of the JUN regulatory mechanism, leading to a high regulon activity of JUN. The JUN regulon specifically accelerates the proliferation and oncogenesis in DSLL, leading to a more terminal differentiatial stage of DSLL (state-2). As a result, state-1 includes cells undergoing rapid cell proliferation and differentiation, transitioning from normal B cells to matured DSLL. In short, DeepMAPS can construct gene regulatory networks and identify cell-type-specific regulatory patterns to offer a better understanding of cell states and developmental orders in diseased subpopulations.

### DeepMAPS provides a multi-functional and user-friendly web portal for analyzing scMulti-omics data

Researchers who lack sufficient computational skills prefer to use webservers or dockers to lessen the programming burden of data analysis. Hence, a code-free and interactive platform for single-cell sequencing data analysis is urgently needed in the public domain. Because of the complexity of single-cell sequencing data, more and more webservers and dockers have been developed in the past three years^53–65^ (**Supplementary Data 11**). However, most of these tools only provide minimal functions such as cell clustering and differential gene analysis. They do not support the joint analysis of scMulti-omics data and especially lack sufficient support for biological network inference. To this end, we developed DeepMAPS as a first-of-its-kind web portal to support online and code-free computational analysis for scMulti-omics data (**Fig. 6a**). The webserver supports the analysis of multiple RNA-seq data, CITE-seq data, and scRNA-ATAC-seq data using DeepMAPS (**Fig. 6b**). Some other methods, e.g., Seurat, are also incorporated as an alternative use for the users’ convenience. Three major steps—data preprocessing, cell clustering and annotation, and network construction—are included in the server. In addition, the DeepMAPS server supports real-time computing and interactive graph representations. Users may register for an account to have their own workspace to store and share analytical results.

**Fig. 6.**
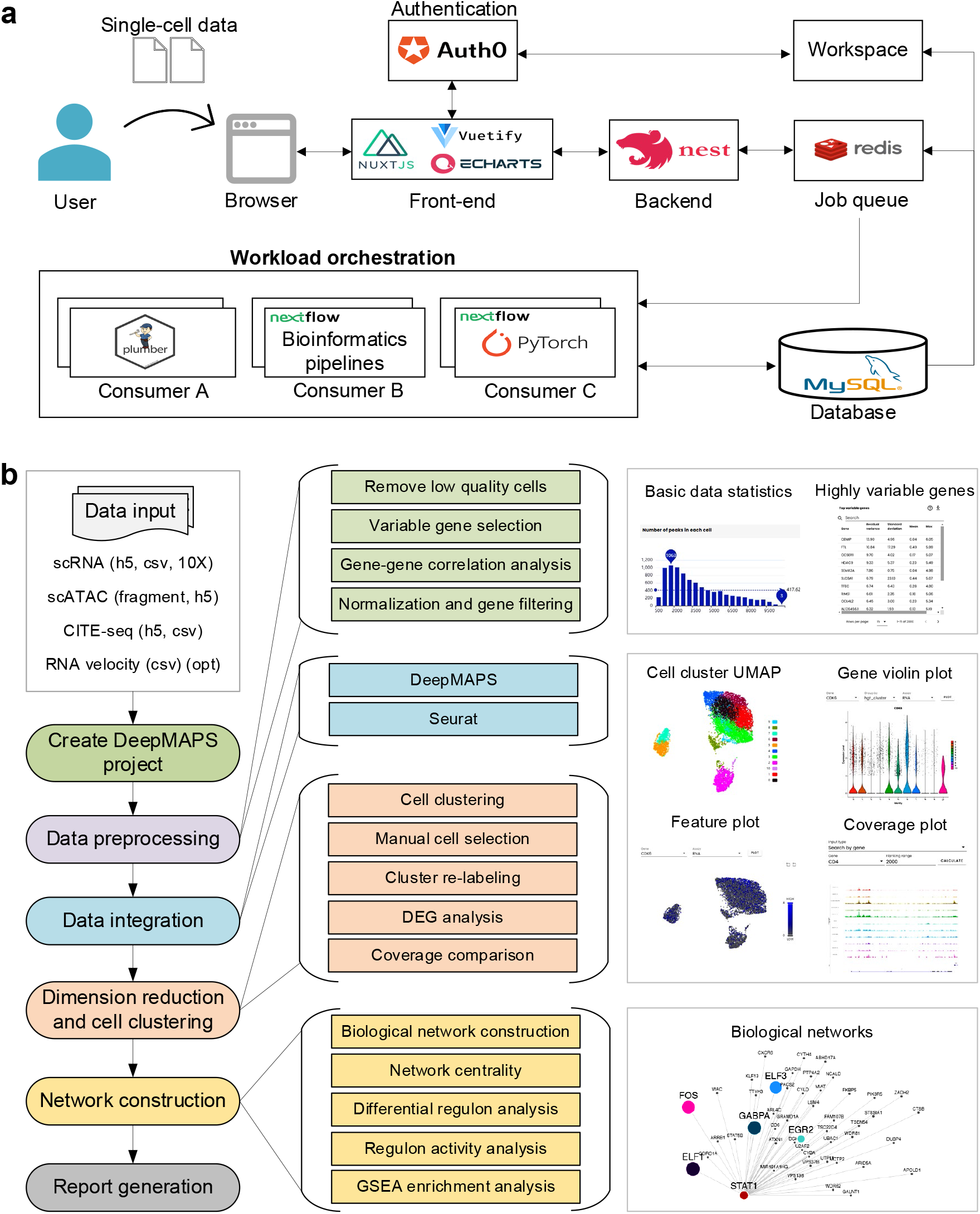
The organization of the DeepMAPS web portal. (a) The software engineering of DeepMAPS and an overview of the framework. (b) Pipeline illustration of the server, including major steps (left), detailed analyses (middle), and featured figures and tables (right).

## Conclusion and discussion

We have highlighted DeepMAPS as the first deep-learning framework that implements heterogeneous graph representation learning and a graph transformer in the study of scMulti-omics data. By building a heterogeneous graph containing both cells and genes, DeepMAPS identifies the joint embedding of them simultaneously and enables the inference of cell-type-specific biological networks along with cell types in an intact framework. Furthermore, the application of heterogeneous graph transformer models the cell-gene relation in an interpretable uniform multi-relations. In such a way, the training and learning process in a graph can be largely shortened to consider cell impacts from a further distance.

By jointly analyzing gene expression and protein abundance, DeepMAPS accurately identified and annotated 13 cell types in a mixed CITE-seq data of PBMC and lung tumor leukocytes based on curated markers that cannot be fully elucidated using a single modality. We also proved that the embedding features identified in DeepMAPS capture statistically significant signals and amplify them when the original signals are noisy. Additionally, we identified biologically meaningful cell-cell communication pathways between DC and TRM CD4^+^ T cells based on the gene network inferred in the two clusters. For scRNA-ATAC-seq, we employed an RNA velocity-based method to dynamically integrate gene expressions and chromatin accessibility that enhanced the prediction of cell clusters. Using such a method, we identified distinct gene regulatory patterns among normal B cells and two DSLL development states. We further elucidated the deep biological connections between cell-cell communications and the downstream gene regulatory networks, which helped characterize and define DSLL states. The identified TFs and genes can be potential markers for further validation and immuno-therapeutical targets in DSLL treatment.

While there are advantages and improved performances for analyzing scMulti-omics data, there is still room to improve the power of DeepMAPS further. First of all, the computational efficiency for super-large datasets (e.g., more than 1 million cells) might be a practical issue considering the complexity of the heterogeneous graph representation (may contain billions of edges). Moreover, DeepMAPS is recommended to be run on GPUs, which leads to a potential problem of reproducibility. Different GPU models have different floating-point numbers that may influence the precision of loss functions during the training process. That is to say, for different GPU models, DeepMAPS may generate slightly different cell clustering and network results, which is the main reason for the development of the webserver. Lastly, the current version of DeepMAPS is based on a bipartite heterogeneous graph with genes and cells. Separate preprocessing and integration steps are required to transfer different modalities to genes for integration into a unique cell-gene matrix. To fully achieve an end-to-end framework for scMulti-omics analysis, the bipartite graph can be extended to a multipartite graph, where each modality can be included as a node type. Such a multipartite heterogeneous graph can also include knowledge-based biological information, such as known molecular regulations and more than two modalities in one graph. However, by including more node types, the computational burden will be increased geometrically, which requires a dedicated discovery of model optimization in the future.

In summary, we evaluated DeepMAPS as a pioneer study for the joint analysis of scMulti-omics data and cell-type-specific biological network inference. It will likely provide new visions of deep learning deployment in single-cell biology. With the development and maintenance of the DeepMAPS webserver, our long-term goal is to create a deep learning-based eco-community for archiving, analyzing, visualizing, and disseminating AI-ready scMulti-omics data.

## Methods

### Data preprocessing and data integration

Different data preprocessing and integration methods are designed for multiple scRNA-seq data, CITE-seq data, and scRNA-ATAC-seq. The method for each type of multi-omic data are shown as follows.

#### Multiple scRNA-seq data

DeepMAPS takes raw scRNA-seq gene expression profiles as inputs. Genes expressed in less than 0.1% of total cells or cells with a maximum of 0.1% of genes expressed are removed. To integrate multiple scRNA-seq datasets, we first reduced the dimension of multiple scRNA-seq data using canonical correlation analysis (CCA) and subsequently searched for mutual nearest neighbors (MNNs) in the shared low-dimensional space. The program then calculates vectors of each cell and corrects gene expressions in each dataset. The output would be an integrated matrix with *I* genes and *J* cells, aggregated from all datasets. Finally, the shared genes were normalized and re-scaled as in Seurat v3. The expression level of gene *i* in cell *j* was denoted as *x_ij_*.

#### CITE-seq data

Low-quality genes and cells are removed as described above. Log normalization is then applied on the RNA matrix *X^R^′* with *I*_1_ genes and *J* cells as follows:

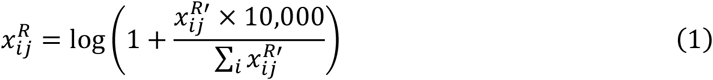

where 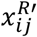 represents the expression of gene *i* in cell *j, i* = 1, 2,…, *I*_1′_, *j* = 1,…, *J*, and 10,000 is the scaling factor. Then, the top 2,000 (or *I_1_* whichever is smaller) highly variable genes are selected from matrix *X^R^*. We still use *X^R^* to represent the submatrix only containing these top-ranked genes.

Subsequently, RNA matrix *X^R^* were spliced by rows with the protein matrix including *I*_2_ proteins to obtain matrix *X′*. Then, a joint centered log-ratio (CLR) normalization was perfomed to obtain the processed matrix *X* as follows:

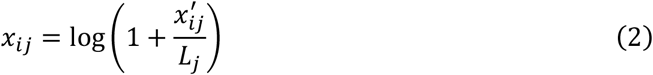

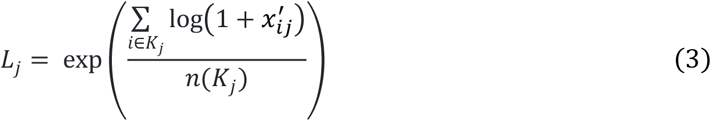

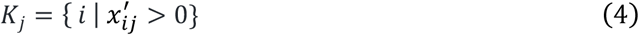

where 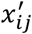 represents the expression for gene/protein *i* in cell *j, i* = 1,…, *min*(2000, *I*_1_) + *I*_2_, *j* = 1,…, *J* and *n*(*K_j_*) represents the number of genes/proteins with non-zero expression values in cell *j*.

#### Matched scRNA-seq and scATAC-seq data (scRNA-ATAC-seq)

Data filtering and quality control are performed to obtain the RNA matrix and binarized ATAC matrix, *X^R^ and X^C^*, respectively, following a standard protocol^5^. We first annotated peak regions in the scATAC-seq based on the method described in MAESTRO^66^. The regulatory potential *R_ik_* for peak *k* and gene *i* is calculated independently by the exponential weight decay with the distance from the peak to the transcription start site (TSS):

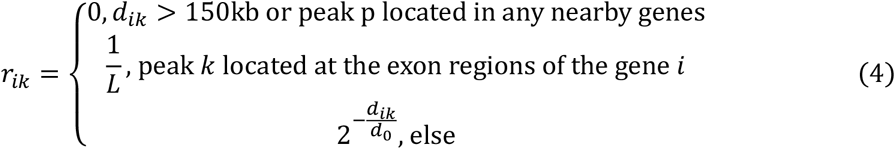

where *d_ik_* is the distance between the center of peak *k* and the TSS of gene *i* and *L* refers to the length of the exon region. To ensure computational efficiency, we set the *R_ik_* as 0 if *d_ik_* was over 150 kb. The default value of *d*_0_ is set to 10kb.

If a peak *k* was located at the exon regions of the gene *i*, the regulatory potential *R_ik_* would be divided by *L*. The gene regulatory activity matrix *X^A^* is defined in which the activity of gene *i* in cell *j* is:

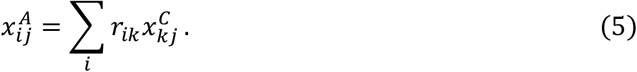

We assume that the activity of a gene to a cell is determined by both gene expression activity and gene regulatory activity with different contributions. Unlike the contribution weights determined directly based on the expression and chromatin accessibility values in Seurat v4 (weighted nearest neighbor)^5^, we hypothesize that the relative contribution of the expression and chromatin accessibility of a gene to a cell is dynamic rather than static and not accurately determined with a snapshot of the cell. RNA velocity is determined by the abundance of unspliced and spliced mRNA in a cell. The amount of unspliced mRNA is determined by gene regulation and gene transcription rate, and the amount of spliced mRNA is determined by the difference between unsliced mRNA and degraded mRNA. We reasoned that for genes with positive RNA velocities, there are higher potentials to drive genes to be transcribed. Thus, its regulatory activity related to chromatin accessibility has a greater influence than the gene expression in defining the overall gene activity in a cell of the current snapshot. For genes with negative velocities, the transcription rate tends to be decelerated, and regulatory activity has less influence on the cell than gene expression activity. We defined a gene activity score (GAS) matrix *X*, of which the score of gene *i* in cell *j* is defined which integrated RNA and ATAC information as follows,

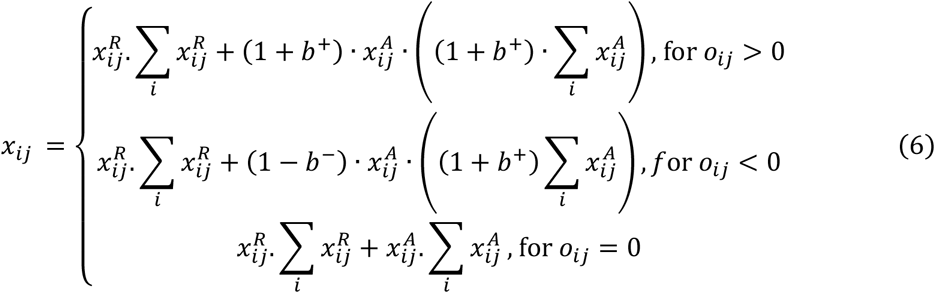

where *x_ij_* and 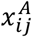 represent the gene expression and gene regulatory activity for gene *i* in cell *j*, respectively, which are devided by the sum of genes, and *o_ij_* denotes the RNA velocity for gene *i* in cell *j*, calculated using CellRank^67^. The weight was defined as:

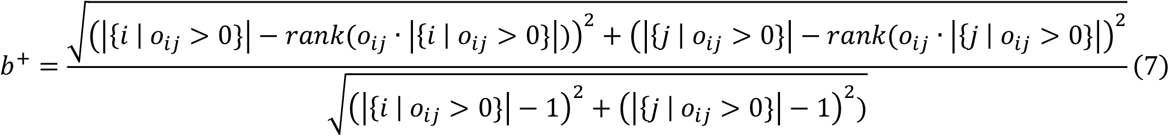

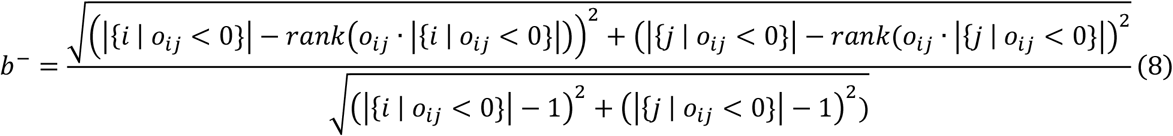

### Construction of gene-cell heterogeneous graph

After the data preprocessing and integrations, we obtaind a matrix, *X* = (*x_ij_*), *i* = 1, 2,…, *I* and *j* = 1, 2,…, *J*, for any types of scMulti-omics data that integrated information from both modalities, with *J* columns for cells and *I* rows for genes. Values in the integrated matrix, *x_ij_*, represent either normalized gene expressions (for multiple scRNA-seq and CITE-seq) or GAS (for scRNA-ATAC-seq). Given an integrated matrix 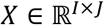 with *I* genes and *J* cells generated in the last step, a heterogeneous graph, denoted as *G* = (*V^C^, V^G^, E*), where 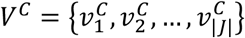 and 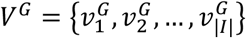 represent the set of cells and genes in the integrated data and *E* = {*e_ij_*}, *i* ∈ *V^G^, j* ∈ *V^C^* represent the edges between gene *i* and cell *j* with the weight corresponding to the expression level of gene *i* in cell *j*. To initialize embedding of each node for gene *i* and cell *j* in the matrix *X*, a feature selection using autoencoder with two layers in both the encoder and decoder was used with the objective function of minimizing the mean squared error (MSE) between *X* and *X′*. The initial embedding of cells and genes are separately trained using the same feature autoencoder architecture with different loss functions. The loss function for gene embeddings is

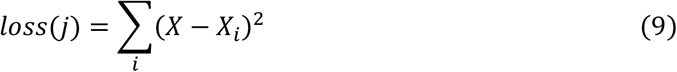

and the loss function for cell embeddings is

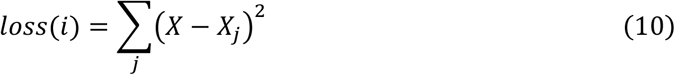

### Learning joint embeddings via a heterogeneous graph transformer

We proposed an unsupervised HGT framework^14^, ^15^ to learn joint embeddings of gene *i* and cell *j*. Given a heterogeneous bipartite graph *G* = (*V^C^, V^G^, E*), DeepMAPS would extracts all linked gene-cell node pairs, which are denoted as (*s, t*), where *t* is the target node and s is the neighbor node of *t*. The training processes consist of four steps: (*i*) calculating multi-head attention; (ii) passing heterogeneous message; (*iii*) aggregating neighbors’ information; and (*iv*) calculating loss function. To learn joint embeddings of gene *i* and cell *j*, we aggregated information from s to get a contextualized representation for *t* and to simultaneously learn the representations of *V_C_* and *V_G_*. It is noteworthy that, to handle the heterogeneous relations in the graph, attention will be calculated via multiple heads where each node type (either cell node or gene node) has a unique head in attention and the nodes are linearly projected to a low dimensional space to model the distribution differences maximally

#### 1) Calculating multi-head attention

We extracted all linked node pairs, where a target node *v_t_* ∈ *V^C^*∪*V^G^* was directly linked to its source node (neighbor node) *v_s_* = *V^C^*∪*V^G^* through edge *e*. Let 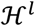 denote the embedding of the *l^th^* HGT layer where *l* ≥ 1, the contextualized representation of *v_t_* on the *l^th^* layer is denoted as 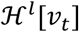, which can be learned by its own presentation layer 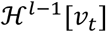 and its neighbor layer 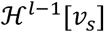 from the (*l* – 1) *^th^* layer. The overall model is formulated as:

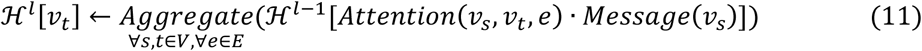

where *Attention* estimates the importance of each neighbor; *Message* extracts the information passed from the neighbors; *Aggregate* is the final step to aggregate the neighborhood message by the attention weight.

The multi-head attention is proposed in the attention level to calculate *h*-head attention for each edge *e* = (*v_s_, v_t_*). Each target node *v_t_* in the *h*-th head, *h* = 1,…, *H*, was mapped into a target-node vector 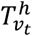 in each HGT layer via linear projection 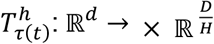, where *D* is the dimension of initial node feature, and 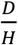 is the feature dimension per head. Similarly, each neighbor node *v_s_* in the head *h* was mapped into a key vector *K^h^*(*v_s_*) with a linear projection 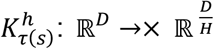:

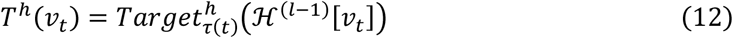

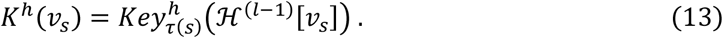

The similarity between the queries and keys was measured (e.g., scaled dot product operator) as attention. Then we calculated the multi-head attention for source node s to target node *t* by the dot product. To maximize parameter sharing while still maintaining the specific characteristics of different relations, we parameterize weight matrices 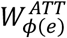 of the interaction operators. The *h^th^* head attention can be defined as:

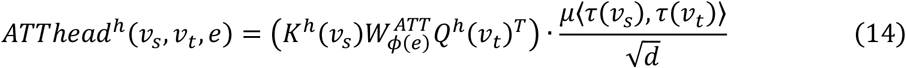

where (·)*^T^* is the transposal function and *μ* is a prior tensor of the significance for each edge *e*, serving as an adaptive scaling to the attention. The attention score in the *h*-th head in the *l^th^* layer was defined as:

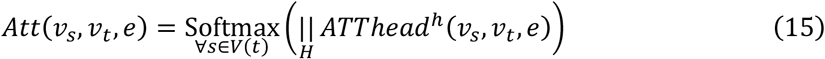

where 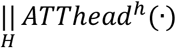 is the concatenation of embeddings in all *H* heads.

#### 2) Passing heterogeneous message

To alleviate the distribution differences of different types of nodes and edges, we incorporated the types of edges into the message passing. The *h^th^* head message for each edge (*v_s_, v_t_*) was defined as:

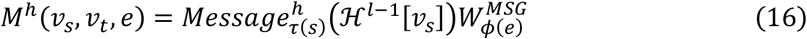

Which means each source node *s* in the head *h* was mapped into a message vector by a linear projection 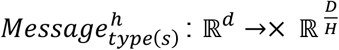, where 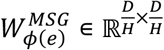 is a distinct edge-based matrix for each edge, and *ϕ*(*e*) is the edge type of the heterogeneous graph. After multi-head aggregation, the degree of message passing was calculated as

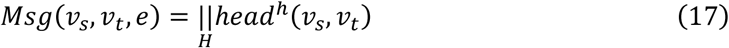

#### 3) Aggregating neighbors’ information

To obtain the representations of each node, we aggregated multi-head attentions and messages. The attention vectors was regarded as the weights for message representation. The representation of target nodes 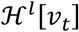 was updated as:

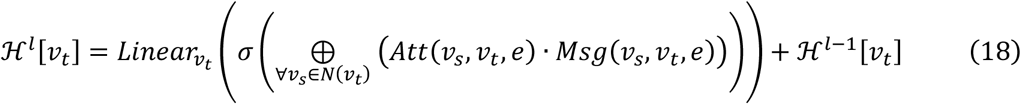

Where *Linear_vt_* is a linear projection.

#### 4) Calculating loss function

The original application of HGT was to solve node classification problems. To utilize HGT for training cell and gene embeddings without supervised classification labels, we used a graph autoencoder (GAE) framework. The whole HGT structure was deployed as an encoder in the GAE, while two embedding matrices *M^C^* and *M^G^* are used to record the trained embeddings of cells and genes from the HGT encoder where the columns were the same embedding dimensions and rows were cells and genes. A decoder was used to reconstruct the heterogeneous graph by the inner product of *M^C^* and *M^G^*. The loss function of GAE was calculated as:

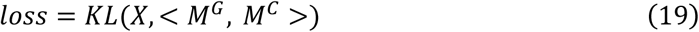

where *KL*(·) represents KL-divergence, and <·> denotes the inner product.

### HGT training on subgraphs

To improve the efficiency and capability HGT model on such a giant heterogeneous graph (tens of thousands of nodes and millions of edges), we performed model training on subgraphs and multiple mini-batches based on the premise of HGSampling^14^. For a sampled node, we added all its direct neighbors into the corresponding budget and added its normalized degree to these neighbors, which were then used to calculate the sampling probability. Such normalization is equivalent to accumulating the probability of random walk of each sampled node to its neighborhood, avoiding the sampling being dominated by high-degree nodes. The sampler constructed a number of small subgraphs from the given giant graph, and these subgraphs were fed into the HGT model in batches with multiple GPUs. Finally, these distributed training results were collected to build the whole graph with representations on the head node.

The core of HGSampling is to sample heterogeneous subgraphs with similar proportions in different types of nodes to avoid sampling highly imbalanced subgraphs in the training process. We sampled a number of subgraphs for each epoch training, and the one-hot of each node was put into the trained model to obtain all node embedding. Given a node *v_t_* which had been sampled and a dictionary *Dict*[*τ*] for each node type *τ*, we added all the first-neighbor nodes of *t* into the corresponding *Dict*[*τ*] and added the normalized degree of *v_t_* to these neighbors to calculate the sampling probability:

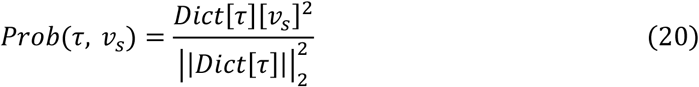

where *Prob*(*τ, s*) is the sampling probability for each source node s of type *τ, Dict*[*τ*] is all node for type *τ* with the normalized degree, ║·║_2_ is the L2-norm, and *Dict*[*τ*][s] is the normalizer degree for the source node s of node type *τ*. Then we sampled all nodes types according to the probability in *Dict*[*τ*], and moved them out of *Dict*[*τ*]. We repeated this sampling 50 times to obtain 50 subgraphs that maximize the coverage of the whole heterogeneous graph, and each subgraph was trained with 100 epochs.

### Cell clustering and cell-type-active gene association network prediction

#### Cell clustering

We apply the Louvain clustering method (igraph v1.2.7, R package) to predict *m* cell clusters {*C_k_, k* = 1, 2,…, *m*} on cell-embedding matrix *M^C^*.

#### Attention-based gene module detection

To infer the connection of genes and cell clusters, we extracted the attention value of gene *i* to cell *j* in the *h*-th head through the step of multi-head attention calculation. We defined the importance function *Imp* of gene *i* to cell *j* as:

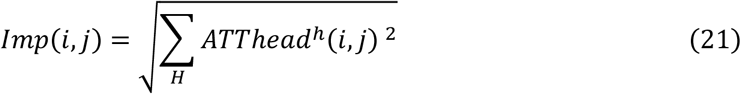

We assigned genes to each cell with a threshold of *mean_g_*(*lmp*(*i, j*)) + *sd_g_*(*Imp*(*i, j*)). A gene is considered to be one of the active genes in a cell, if *Imp* is higher than the threshold.

#### The Steiner Forest Problem (SFP) model

We build an SFP model on a heterogeneous graph to extract the most critical gene-gene and gene-cell relations contributing to the gene module specificity in a cell cluster. The input of this model includes three parts:

1. gene-gene relations 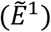 defined by the embedding (resulted from HGT) Pearson’s correlation between genes,
2. gene-cell relations are defined by the attention score of a gene to a cell 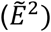.
3. a set of cell clusters, {C_k_, k = 1, 2,…, m}, predicted by the HGT model.

We define a weighted heterogeneous graph 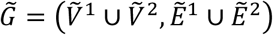 in which nodes represent genes 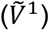 and cells 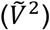, and edges represent both gene-gene 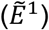 and gene-cell 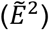 relations. We formulate this problem using a combinatorial optimization model defined as below

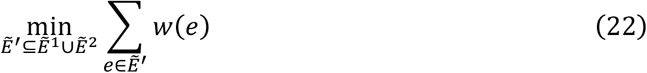

s.t.

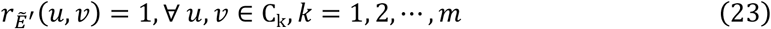

where 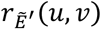 is a binary indicator function representing whether two nodes, *u* and *v*, are connected (1) or not (0) in the subgraph induced by 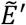 in 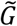. We aim to identify the minimum weighted edge set, 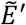, from the heterogeneous graph 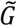, so that cells in the same cell type could be connected to each other via edges in 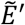.

First, given the huge size of 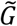, containing 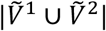 nodes and 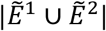 edges, we convert 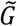 into a sparser graph, 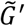, by iteratively finding a global alignment between genes and cells based on the gene-cell edges, using the maximum matching theory^68^. In graph theory, a matching or independent edge set in an undirected graph is a set of edges without common nodes, and a maximum matching in a weighted graph is a matching 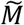 that yields the maximum sum of edge weights. To fulfill this task, we build a weighted bipartite graph, 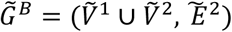, by only retaining the gene-cell edges. The objective is to identify an edge subset, 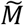, to satisfy

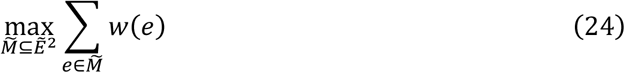

We calculate 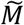 of 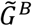 using the igraph R package^69^. Then, we remove the cell nodes incident by edges in *M*. The prediction of maximum matching and deletion of cell nodes incident to edges in previously identified maximum matchings is repeated until there is no cell node in the remaining graph. We compute the union of all the matchings as 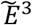, and then construct 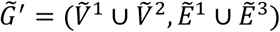. Finally, the weights of gene-cell edges, 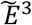, and gene-gene edges, 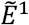, are normalized by the following two functions, respectively.

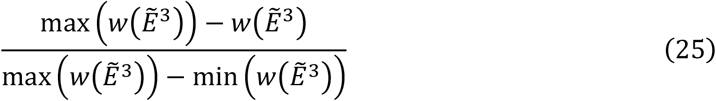

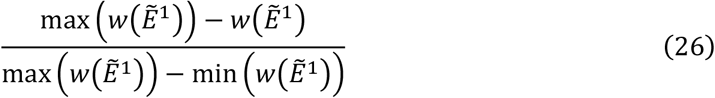

Second, we find the edge set, 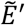, of the SFP, 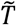, as follows. To begin, we utilize the igraph R package to calculate a minimum spanning forest (MSF)^68^, 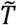, of 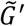. An MSF means that each pair of nodes in the same connected component could be connected to each other. Herein, we only need the edges to connect cell nodes belonging to the same cell type. Therefore, we iteratively remove the gene nodes with degree one and their unique neighbor is not a cell node from 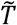, until no such gene node exists in 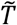. Finally, we output the edge set of 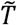, i.e., 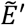, as the solution to the SFP model.

For each connected component of the SFP, the gene-gene edges denote the co-expression relations among genes in the same module, while the set of gene-cell edges represents the cell type specificity of this gene module, and this module is a cell-type-active gene module.

### Construct gene regulatory network from scRNA-ATAC-seq data

#### Infer master TFs and GRNs in each cell type

To quantify the intensity of genes regulated by TFs, we design a regulatory intensive (RI) score, which can be decomposed into two components: (i) the regulatory potential (*R_ip_*) of peaks calculated in the preprocessing step, and (ii) the binding affinity (BA) score of TFs to the peaks. The TF binding profiles were obtained from the JASPAR database. To reduce false positives of binding sites, we selected significant binding sites with *p*-values for TF binding profile matches less than 0.0001. The BA score was the transformed relative score obtained from TF binding profiles, and the RI score of TF *q* to the gene *i* in the cell *j* was then defined as:

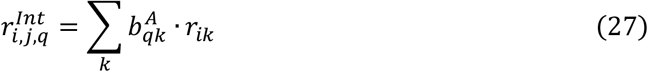

Master TFs are genes at the top of a gene regulation hierarchy, particularly in regulatory pathways related to cell fate and differentiation. To infer cell type master TFs, we constructed cell-type-specific GRN with an RI score as edge weight and calculate centrality, which reflected the importance of each node in the network to rank the TFs in each cell type. TFs with high ranks were regarded as master TFs. Considering the RI score of TFs to genes, eigenvector centrality which assigned relative scores to all nodes in the network based on the concept that connections to high-scoring nodes contributed more to the score of the node in question than equal connections to low-scoring nodes is applied to infer master TFs. The eigenvector centrality (*EC_v_*) of a node *v* in GRN was defined as:

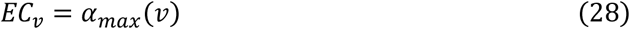

where *a_max_* is the eigenvector corresponding to the largest eigenvalue of the weighted adjacency matrix of a GRN.

#### Identity differential regulon (CTSRs)

We identify CTSRs by logFC and Wilcoxon rank-sum test to detect regulons associated with disease states. For a cell type active regulon, we defined a regulon activity score (RAS) as:

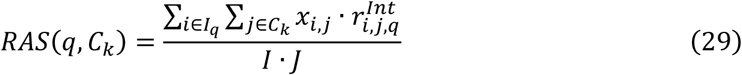

where *I_q_* denotes genes regulated by TF *q* in cell type *C_k_*. The significance of the difference is calculated using the Wilcoxon rank-sum test. If the BH-adjusted *p*-value is less than 0.05 between different cell clusters and the log fold change larger than 0.10, we consider the regulon to be differentially active in this cluster, and it is defined as a CTSR.

### Benchmarking quantification and statistics

#### Adjusted rand index (ARI)

ARI is used to compute similarities by considering all pairs of the samples assigned in clusters in the current and previous clustering adjusted by random permutation. A contingency table is built to summarize the overlaps between the two cell label lists with *b* elements (cells) to calculate the ARI. Each entry denotes the number of objects in common between the two label lists. The *ARI* score can be calculated as:

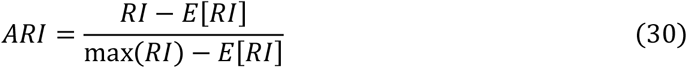

Where *E*[·] is the expectation, *RI* is the unadjusted rand index, which is defined as,

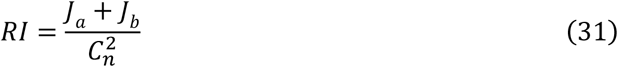

where *J_a_* is the number of cells that are assigned to the same cell cluster as benchmark labels, and *J_b_* is the number of cells that are assigned to different cell clusters as benchmark labels.

#### Average Silhouette Width (ASW)

Unlike ARI, which requires known ground truth labels, silhouette weight refers to a method of interpretation and validation of consistency within clusters of data. The silhouette weight indicates how similar an object is to its cluster (cohesion) compared to other clusters (separation). The silhouette width ranges from −1 to +1, where a high value indicates that the object is well matched to its cluster. The silhouette score *sil*(*j*) can be calculated by:

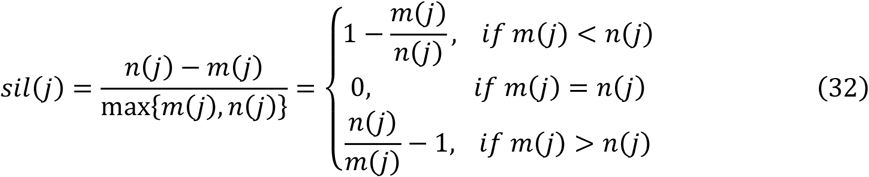

where *m*(*j*) is the average distance between a cell *i* and all other cells in the same cluster, and *n*(*j*) is the average distance of *i* to all cells in the nearest cluster to which *j* does not belong. We take the average silhouette of all cells in a cluster as the average silhouette weight (ASW) to represent the cell distance of the cell cluster.

#### Closeness centrality (CC)

The closeness centrality (CC)^70^ of a vertex *u* is defined by the inverse of the sum length of the shortest paths to all the other vertices *v* in the undirected weighted graph. The formulation is defined as:

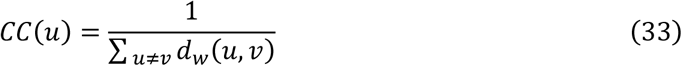

where *d_w_*(*u, v*) is the shortest weighted path between u and v. If there is no path between vertex u and v, the total number of vertices is used in the formula instead of the path length. The CC is calculated using igraph R package with function igraph::betweenness.

#### Eigenvector centrality (EC)

Eigenvector centrality (EC)^71^ scores correspond to the values of the first eigenvector of the graph adjacency matrix. The EC score of vertex *u* is defined as:

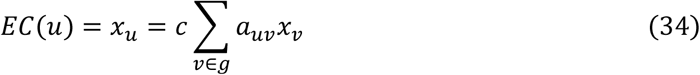

where *c* is inverse of the eigenvalues of eigenvector *x* = [*x*_1_, *x*_2_,…, *x_n_*], *a_uv_* is the adjacent weighted matrix of undirect graph *g*. The EC is calculated using igraph R package with function igraph::evcent.

#### Pathway enrichment test

To evaluate the function of the regulatory network, we use pathway enrichment analysis^72^ to identify pathways that are significantly represented in each cell cluster active regulon and count the number of regulon-enriched pathways. The pathway enrichment analysis is performed using enrichR R package ^73^.

### Robustness evaluation

#### Cell cluster leave-out test

For a benchmark dataset with a real cell type label, we removed all cells in one cell type and ran DeepMAPS. Then, we traverse all cell types (one at a time) to evaluate the robustness with ARI. We removed cells in predicted cell clusters from DeepMAPS and other benchmark tools for data without benchmark labels.

#### Read depth simulation test

We performed a down-sampling simulation for gene expressions to test the robustness of DeepMAPS to read depth. Let matrix *C* be the *N* × *M* expression count matrix, where *N* is the number of cells and *M* is the number of genes. Define the cell sequencing depths 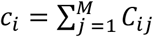, i.e., the column sums of *C*. Thus, the average sequencing depth of the experiment is 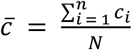. Let 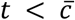 be our target downsampled sequencing depth and let *C*\* be the *N* × *M* downsampled matrix. We perform the downsampling as follows:

For each spot *i* = 1,…, *N*:

1. Define the total counts to be sampled in the cell *i* as 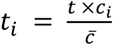
2. Construct the character vector of genes to be sampled as 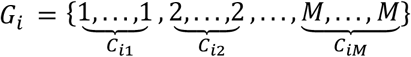
3. Sample *t_i_* elements from *G_i_*, without replacement and define *N_j_* as the number of times gene *j* was sampled from *G_j_*, for *j* = 1,…, *M*.
4. Let 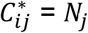

Using this method, the average downsampled sequencing depth is:

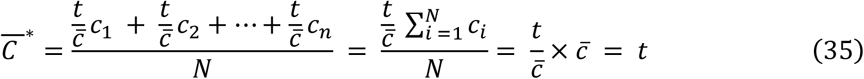

Note that, this method preserves the relative total counts of each cell, i.e., cells that had higher sequencing depths in the original matrix have proportionally higher depths in the downsampled matrix.

### Comparisons with existing tools

In order to assess the performance of DeepMAPS alongside other proposed scMulti-omics benchmark tools, we compared DeepMAPS with Seurat (v 3.2.3 and v 4.0.0, https://github.com/satijalab/seurat), MOFA+ (v 1.0.0, https://github.com/bioFAM/MOFA2), Harmony (v 0.1, https://github.com/immunogenomics/harmony), and TotalVI (v 0.10.0, https://github.com/YosefLab/scvi-tools). Because of the integration capability, DeepMAPS was compared with Seurat v 3.2.3 and Harmony on multiple scRNA-seq data, with Seurat v4.0.0, MOFA+, and TotalVI on CITE-seq data, and with Seurat v4.0.0 and MOFA+ on scRNA-ATAC-seq data. All benchmark tools used the default settings. We also evaluated the performance of gene association network inference with IRIS3^11^ and a normal gene co-expression network inference method. Specifically, in IRIS3, cell-gene biclusters were first identified using QUBIC2^74^. Cell-type-active gene modules were identified in each cell cluster (using the same cell label predicted in DeepMAPS to ensure comparability) by performing a cell-wise hypergeometric enrichment.

On the other hand, all genes are selected to calculate a gene expression correlation score (Pearson’s correlation) between any pair of two genes using cells in one cell cluster. Gene pair expression correlations with a BH-adjusted p-value smaller than 0.05 are retained and used to build the overall co-expression network in one cell cluster. Co-expressed sub gene modules are inferred by performing Louvain clustering on the co-expression network. For scRNA-ATAC-seq data, we compared regulon and cell-type-specific regulon inferred from DeepMAPS with IRIS3 in terms of enriched biological pathways. Noted that IRIS3 only supports regulon inference from scRNA-seq data based on *de novo* motif findings; thus, we used the GAS matrix generated in DeepMAPS as an input of IRIS3.

### DeepMAPS server construction

DeepMAPS runs on an HPE XL675d RHEL system with 2×128-core AMD EPYC 7H12 CPU, 64GB RAM, and 2×NVIDIA A100 40GB GPU. The backend is written in TypeScript using the NestJs framework. Auth0 is used as an independent module to provide user authentication and authorization services. Redis houses a queue of all pending analysis jobs. There are two types of jobs in DeepMAPS: The stateful jobs are handled by the Plumber R package to provide real-time interactive analysis; the stateless jobs, such as CPU-bound bioinformatics pipelines and GPU training tasks that could take a very long time, are constructed using Nextflow (**Supplementary Data 12**). All running jobs are orchestrated using Nomad, allowing each job to be assigned with proper cores and storage and keeping jobs scalable based on the server load. The job results are deposited to a MySQL database. The frontend is built with NUXT, Vuetify as the UI library, Apache ECharts, and Cytoscape.js for data visualization. The frontend server and backend server are communicated using REST API.

## Data availability

All data used for benchmarking and case study are collected from the public domain and can be retrieved using links or accession numbers provided in **Supplementary Data 1.**

## Code availability

The source code of DeepMAPS Docker is freely available at (https://github.com/OSU-BMBL/deepmaps). The DeepMAPS webserver is available at https://bmblx.bmi.osumc.edu/).

## Acknowledgments

This work was supported by awards R01-GM131399, R35-GM126985, and U54AG075931 from the National Institute of General Medical Sciences of the National Institutes of Health. The work was also supported by award NSF1945971 from the National Science Foundation. In addition, we thank Dr. Fei He from the Northeast Normal University for his valued suggestions in framework construction and data testing.

## Contributions

Q.M., B.L., and D.X. conceived the basic idea and designed the framework. X.W. wrote the backbone of DeepMAPS. C.W. built the backend and frontend server. S.G. designed the interactive figures on the server. Y. Liu carried out RNA velocity calculations. Y. Li designed the SFP model for gene module predictions. A.M, X.W., and J.L. carried out benchmark experiments. X.W., Y.C., and B.L. performed robustness tests. A.M., J.L., X.W., X.G., and T.X. carried out the case study. J.W., D.W., Y.J., J.L., and L.S. performed tool optimizations. A.M. led the figure design and manuscript writing. All authors participated in the interpretation and writing of the manuscript.

