## Supplementary Fig. 1-5 for "DeepMAPS: Single-cell biological network inference using heterogeneous graph transformer"

[$] To whom correspondence should be addressed

**Supplementary Fig. 1.** UMAPs of all 128 embeddings of the CITE-seq data.

**Supplementary Fig. 2.** Gene associated networks generated in the four clusters based on the CITE-seq case 1 data.

**Supplementary Fig. 3.** Performance comparison of cell clustering in the eight benchmark scRNA-ATAC-seq datasets.

**Supplementary Fig. 4.** Regulon activity heatmap of regulons of normal B cells and DSLL.

**Supplementary Fig. 5.** Gene expression, chromatin accessibility, GAS, and attention score of genes regulated by JUN in normal B cells and DSLL.

**Supplementary Data (separate file)**

**Supplementary Data 1.** Datasets used in this paper for benchmarking and case study.

**Supplementary Data 2.** Grid optimization of DeepMAPS on multiple RNA-seq data.

**Supplementary Data 3.** Grid optimization of DeepMAPS on CITE-seq data.

**Supplementary Data 4.** Grid optimization of DeepMAPS on matched RNA-seq and ATAC-seq data.

**Supplementary Data 5.** Robustness comparison of DeepMAPS on cell cluster leave-out.

**Supplementary Data 6.** Robustness comparison of DeepMAPS on simulated read depth.

**Supplementary Data 7.** Benchmark network results.

**Supplementary Data 8.** CITE-seq marker gene list for each cell cluster.

**Supplementary Data 9.** RNA+ATAC-seq case study marker gene list for each cell cluster.

**Supplementary Data 10.** Regulon scores for the top 20 centrality TF-regulons in each of the three B cell clusters.

**Supplementary Data 11.** Comparison of public single-cell analysis servers.

**Supplementary Data 12.** Computing time comparison.

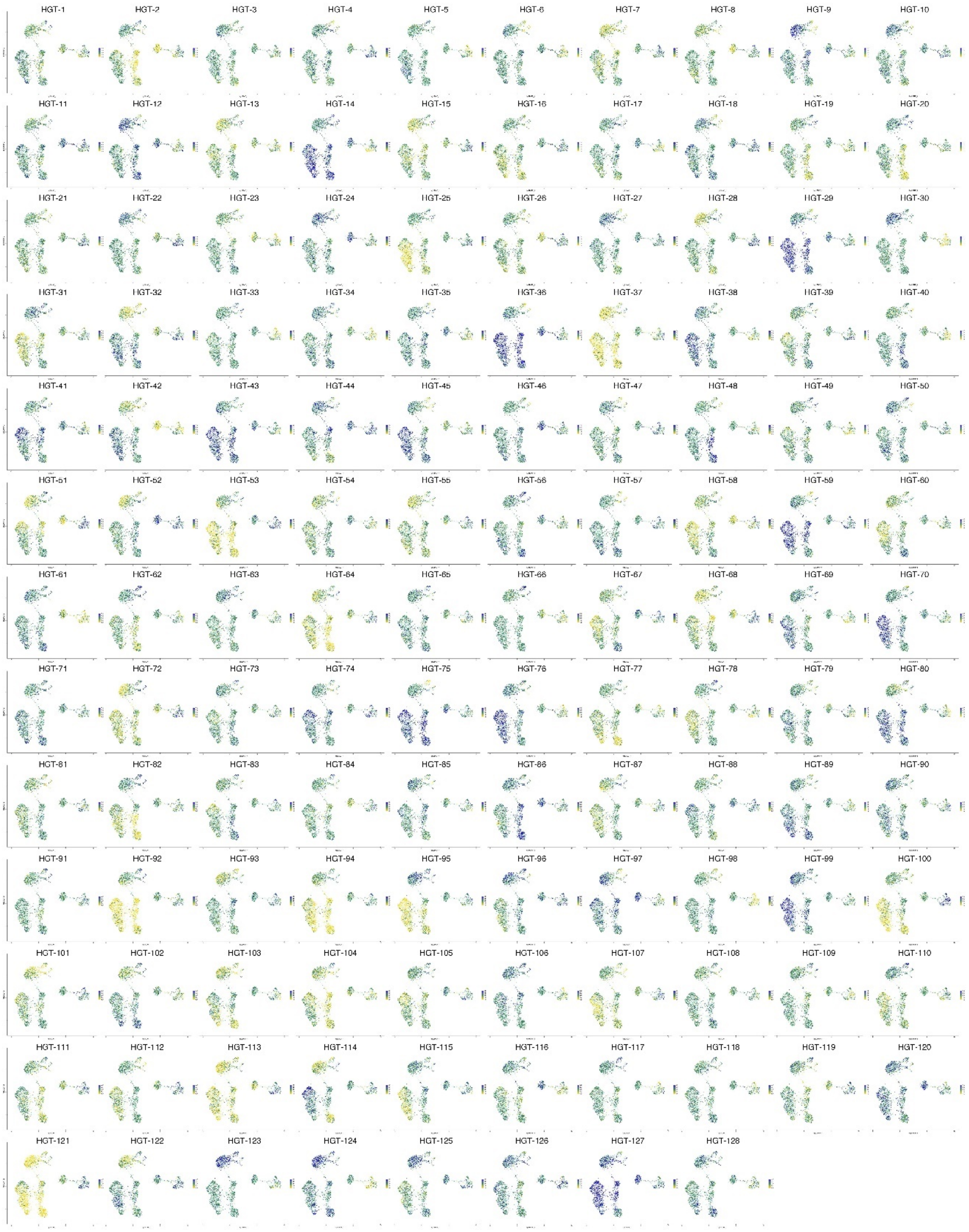


**Supplementary Fig. 1**. UMAPs of all 128 embeddings of the CITE-seq data. Each embedding showed signals that help separate cell clusters. The sequence of embedding is not ranked.

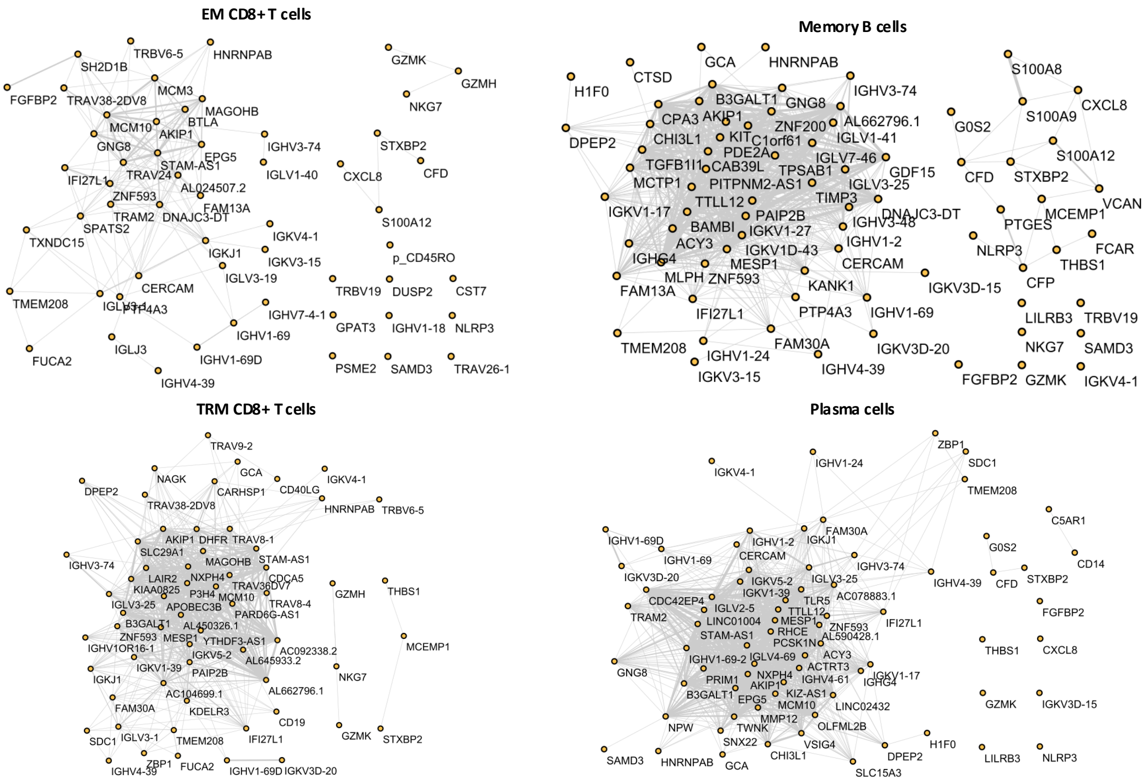


**Supplementary Fig. 2**. Gene associated networks generated in the four clusters based on the CITE-seq case 1 data. Each dot presents a gene with high attention scores in the cluster. Each edge represents the correlation coefficient of the HGT embedding of the two genes—a higher correlation coefficient results in a thicker edge. Only correlation coefficients higher than 0.8 were kept.

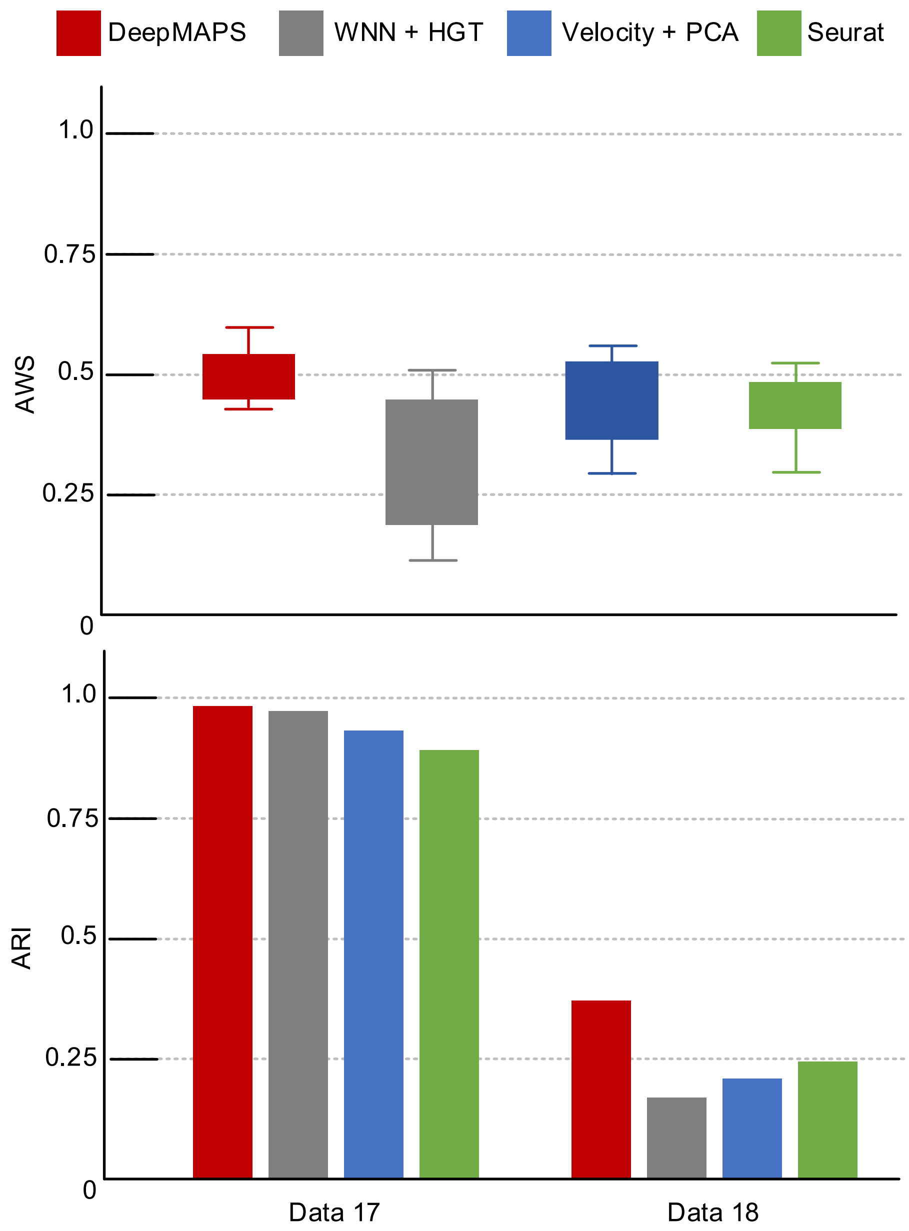


**Supplementary Fig. 3**. Performance comparison of cell clustering in the eight benchmark scRNA-ATAC-seq datasets. We compared DeepMAPS with (1) Seurat WNN + HGT, (2) DeepMAPS velocity-weighted + Seurat PCA, and (3) Seurat (WNN+PCA). For data 11−16 without benchmark labels, we compared AWS as similar as in the benchmarking section; for Data 17−18, we compared the ARI.

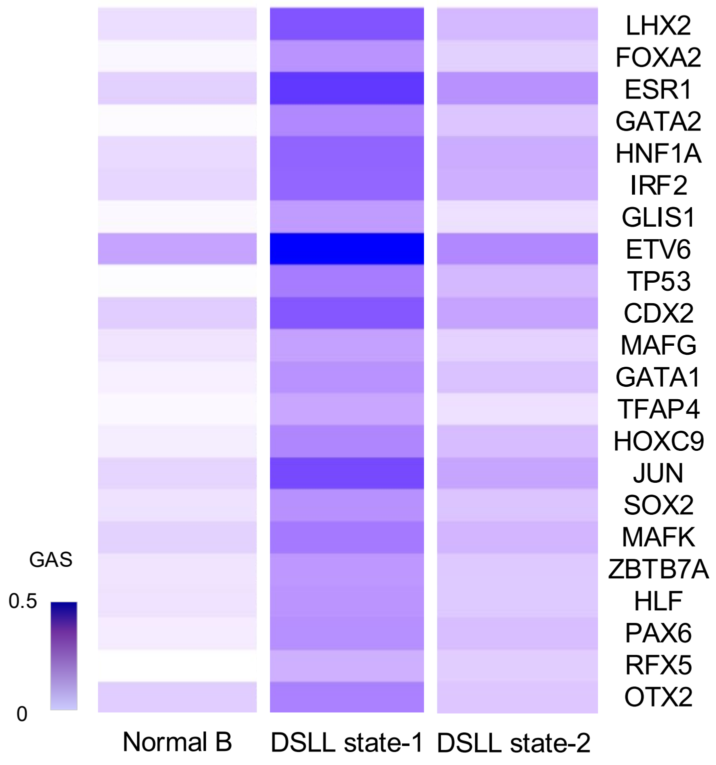


**Supplementary Fig. 4**. Regulon activity heatmap of regulons in DSLL state-1 that are differentially active compared to normal B cells and DSLL state-2.

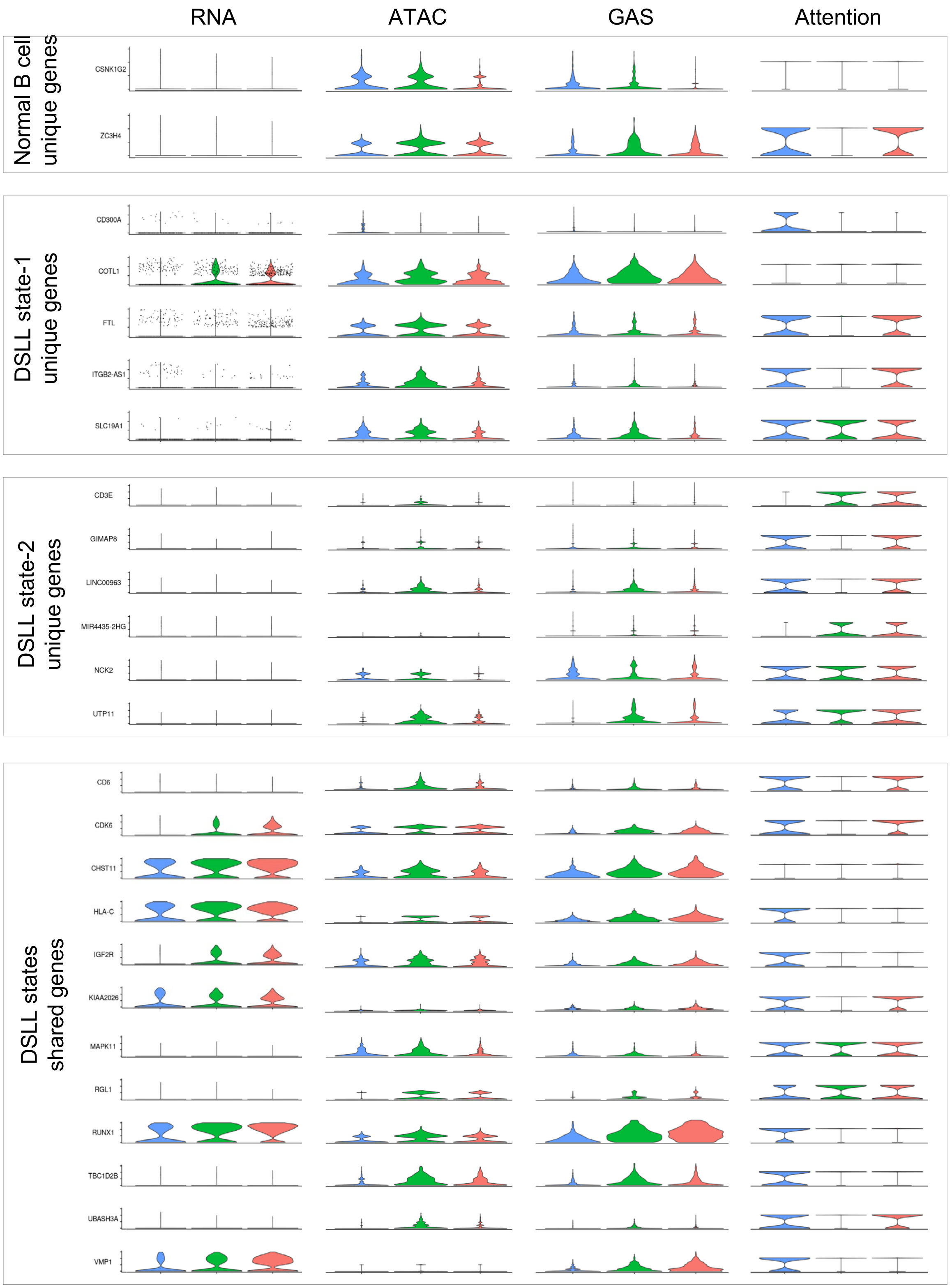


**Supplementary Fig. 5**. Gene expression, chromatin accessibility, GAS, and attention score of genes regulated by JUN in normal B cells (blue), DSLL state-1 (green), and DSLL state-2 (pink).
